# Bacterial surface lipoproteins mediate epithelial microinvasion by *Streptococcus pneumoniae*

**DOI:** 10.1101/2023.10.18.562909

**Authors:** Jia Mun Chan, Elisa Ramos-Sevillano, Modupeh Betts, Holly U. Wilson, Caroline M. Weight, Ambrine Houhou-Ousalah, Gabriele Pollara, Jeremy S. Brown, Robert S. Heyderman

## Abstract

*Streptococcus pneumoniae*, a common coloniser of the upper respiratory tract, invades nasopharyngeal epithelial cells without causing disease in healthy people. We hypothesised that surface expression of pneumococcal lipoproteins, recognised by the innate immune receptor TLR2, mediate epithelial microinvasion. Mutation of *lgt* in serotype 4 (TIGR4) and serotype 6B (BHN418) pneumococcal strains abolishes the ability of the mutants to activate TLR2 signalling. Loss of *lgt* also led to concomitant decrease in interferon signalling triggered by the bacterium. However, only BHN418 *lgt::cm* but not TIGR4 *lgt::cm* was significantly attenuated in epithelial adherence and microinvasion compared to their respective wild-type strains. To test the hypothesis that differential lipoprotein repertoires in TIGR4 and BHN418 lead to the intraspecies variation in epithelial microinvasion, we employed a motif-based genome analysis and identified an additional 525 a.a. lipoprotein (<u>p</u>neumococcal <u>a</u>ccessory lipoprotein <u>A</u>; *palA*) encoded by BHN418 that is absent in TIGR4. The gene encoding *palA* sits within a putative genetic island present in ∼10% of global pneumococcal isolates. While *palA* was enriched in carriage and otitis media pneumococcal strains, neither mutation nor overexpression of the gene encoding this lipoprotein significantly changed microinvasion patterns. In conclusion, mutation of *lgt* attenuates epithelial inflammatory responses during pneumococcal-epithelial interactions, with intraspecies variation in the effect on microinvasion. Differential lipoprotein repertoires encoded by the different strains do not explain these differences in microinvasion. Rather, we postulate that post-translational modifications of lipoproteins may account for the differences in microinvasion.

**IMPORTANCE:** *Streptococcus pneumoniae* (pneumococcus) is an important mucosal pathogen, estimated to cause over 500,000 deaths annually. Nasopharyngeal colonisation is considered a necessary prerequisite for disease, yet many people are transiently and asymptomatically colonised by pneumococci without becoming unwell. It is therefore important to better understand how the colonisation process is controlled at the epithelial surface.

Controlled human infection studies revealed the presence of pneumococci within the epithelium of healthy volunteers (microinvasion). In this study, we focused on the regulation of epithelial microinvasion by pneumococcal lipoproteins. We found that pneumococcal lipoproteins induce epithelial inflammation but that differing lipoprotein repertoires do not significantly impact the magnitude of microinvasion. Our results highlight the potential importance of the post-translational modification of lipoproteins in the mediation of epithelial invasion during pneumococcal colonisation. Targeting mucosal innate immunity and epithelial microinvasion alongside the induction of an adaptive immune response may be effective in preventing pneumococcal colonisation and disease.

## INTRODUCTION

*Streptococcus pneumoniae* (pneumococcus) is a versatile pathobiont capable of asymptomatically colonising the nasopharynx, causing localised infections of the middle ear, respiratory tract and lungs, and causing disseminated invasive disease (e.g. bacteraemic pneumonia and meningitis) with high mortality rates (1). *S. pneumoniae* has long been considered an extracellular pathogen despite demonstration of bacterial invasion *in vitro* using epithelial and endothelial cell lines (1). However, controlled human infection with a serotype 6B strain revealed that the pneumococcus invades the nasopharyngeal epithelium of healthy carriers, stimulating epithelial inflammation without causing overt symptoms or disease (2–4). We have termed this phenomenon microinvasion, which is distinct from the invasion of deeper tissues or dissemination systemically which characterise disease (2). Inflammation triggered by the epithelium-associated and intracellular bacteria, which peaks 9 days post inoculation, may be important for clearance and onward transmission (2).

In this study, we explored the hypothesis that surface expression of pneumococcal lipoproteins mediate epithelial microinvasion. Pneumococcal lipoproteins are post-translationally lipidated surface proteins, many of which function as metabolite transporters (5, 6). *S. pneumoniae* lipoproteins have also been shown to be major TLR2 ligands in macrophages, are required for a Th17 response and for many of the dominant macrophage gene transcriptional responses, such as induction of IRAK-4–dependent protective cytokines (7–9). *S. pneumoniae* encodes over 30 lipoproteins, including the bifunctional adhesin/manganese transporter PsaA and the peptidoglycan hydrolase DacB (5, 10–12). Blocking lipidation by mutating the prolipoprotein diacylglyceryl transferase encoding gene *lgt* de-anchors lipoproteins from the cell surface, resulting in the release of immature preprolipoproteins into the extracellular milieu and abolishing the ability of the bacteria to activate TLR2 signalling (8, 9, 13). Mutating *lgt* also attenuates pneumococcal virulence and shortens colonisation duration in murine models (8, 14).

To explore whether heterogeneity in surface-expression of pneumococcal lipoproteins also explains the differences in microinvasion seen between strains, we blocked lipoprotein lipidation by inactivation of *lgt* in two strains: a highly invasive strain (TIGR4, serotype 4) and a less invasive strain (BHN418, serotype 6B) which was used in the controlled human challenge experiments (15, 16). It is important to note that pneumococcal strains from both serotypes can asymptomatically colonise as well as cause invasive disease in susceptible hosts, albeit to different extents (17). Our findings indicate that lipoproteins contributed to epithelial inflammation by activating the TLR2 pathway and augmenting an interferon response. While attenuation of inflammatory responses were seen with both serotype 6B and serotype 4 *lgt* mutants, we observed intraspecies differences in the contribution of lipoproteins to microinvasion, with greater effects of lipoproteins with the less invasive 6B strain.

Genomic analysis revealed the presence of a previously uncharacterised lipoprotein encoded within a genetic island found in BHN418 and approximately 10% of pneumococcal strains, but not in TIGR4. We designate this protein <u>p</u>neumocccal <u>a</u>ccessory lipoprotein <u>A</u>, or PalA. We hypothesized *that palA* mediate the intraspecies differences in microinvasion, however, cell culture and murine colonisation experiments did not show an essential role for *palA* in microinvasion, colonisation or disease. Our results suggest that the contribution of lipoproteins and lipoprotein processing to microinvasion may be more complex and we propose may occur through differential post-translational lipidation.

## RESULTS

### Pneumococcal *lgt* mutants induce lower levels of TLR2 and interferon signalling compared to than wild type strains

In line with previous reports, mutation of *lgt* in both TIGR4 and BHN418 completely abolished the ability of these strains to trigger TLR2 signalling in HEK-Blue^TM^ hTLR2 reporter cells, while genetic complementation of *lgt* at a chromosomal ectopic site restored WT-like ability to stimulate the TLR2 pathway (Figure 1A) (9). Although macrophages respond to pneumococcal infections by activating TLR2 signalling pathways, it is unknown if nasopharyngeal epithelial cells respond in the same way (8, 9, 18). Using a transcriptional module reflective of TLR2 signalling and previously published transcriptomic datasets (2, 19), we found evidence of elevated TLR2-mediated transcriptional activity in nasopharyngeal epithelial cells infected with TIGR4 and BHN418 (Figure 1B; Supplementary Figure 1).

TLR2 activation is necessary for full induction of TLR4 by the *S. pneumoniae* virulence factor pneumolysin (20, 21). Transcriptomic analyses of human nasal biopsy samples from controlled pneumococcal challenge experiments and nasopharyngeal cell lines infected with *S. pneumoniae* also showed upregulation of interferon signalling (6, 7). We therefore hypothesize that TLR2 activation potentiates interferon signalling in epithelial cells triggered by *S. pneumoniae* infection. Using qPCR, we observed that Detroit 562 cells infected with TIGR4 *lgt::cm* have reduced expression of *CXCL10*, *IFNB1* and *IFNL1* compared to cells infected with WT TIGR4 (Figure 1C-E), while cells infected with BHN418 *lgt::cm* have reduced expression of *CXCL10* and *IFNL3* compared to those infected with WT BHN418 (Figure 1C,1F). Our results suggest that lipoprotein-mediated TLR2 activation augments the epithelial interferon response during pneumococcal microinvasion.

**Figure 1.**
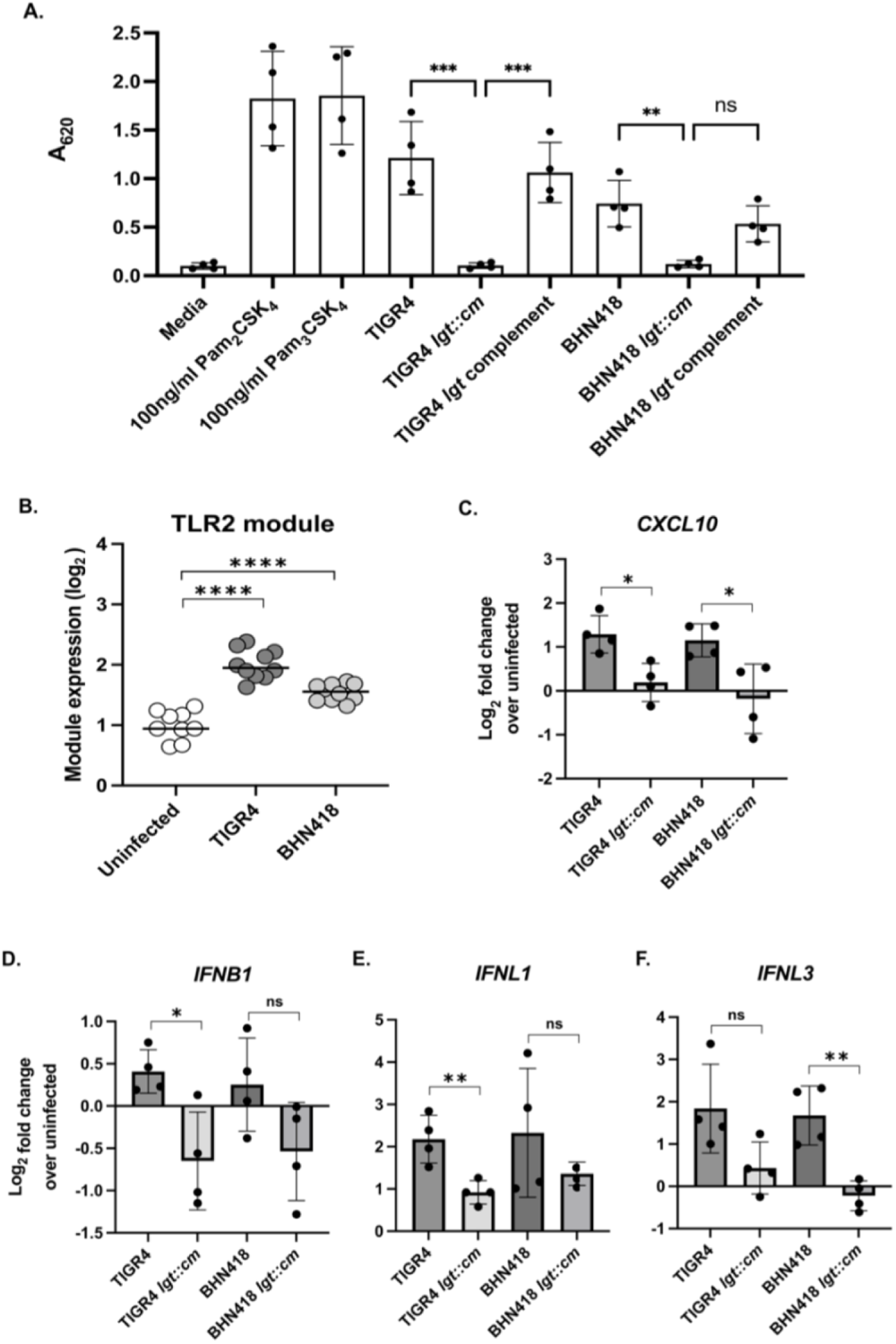
Pneumococcal *lgt* mutants were less inflammatory compared to WT strains. (A) SEAP reporter readout from HEK-Blue^TM^ hTLR2 reporter cells treated with pneumococcal strains at MOI 10 for 16 hours. (B) Expression of a transcriptional module reflective of TLR2-mediated activity in Detroit 562 cells infected with TIGR4 and BHN418 for 3 hours. (C-F) Transcript levels of (C) *CXCL10*, (D) *IFNB1*, (E) *IFNL1* and (F) *IFNL3,* quantified via qPCR using total RNA extracted from Detroit 562 cells after 6 hours of infection with pneumococcal strains. Statistical significance was determined using multiple comparison test with Bonferroni’s correction (A), Mann-Whitney test (B), or Student’s *t-*test assuming equal variance (C-F). * indicates *p* < 0.05, ** indicates *p* < 0.01, *** indicates *p* < 0.001, *** indicates *p* < 0.0001

### .Mutation of *lgt* attenuates epithelial microinvasion by *S. pneumoniae* serotype 6B but not by serotype 4

To determine if mutation of *lgt* and loss of TLR2 signalling impact on pneumococcal microinvasion, we infected confluent Detroit 562 nasopharyngeal cells (NPE) with the serotype 6B (BHN418) and 4 strains (TIGR4) for 3 hours (3 hpi), measuring the number of cell associated, intracellular and planktonic bacteria in the cell culture supernatant. Mutation of *lgt* significantly attenuated the ability of BHN418 but not TIGR4 to associate with and be internalised into Detroit 562 cells (Figure 2A-B, 2D-2E). In concordance with prior reports, serotype 4 strains were more invasive compared to serotype 6B strains, with ∼5 times more intracellular WT TIGR4 recovered compared to WT BHN418 (Figure 2B, 2D) (15, 17). The *lgt* mutation also significantly reduced the number of planktonic BHN418 but not TIGR4 (Figure 2F). Genetic complementation of *lgt* in the BHN418 *lgt::cm* mutant did not fully restore microinvasion of NPE cells to WT-like levels, despite complementation in the HEK-Blue^TM^ hTLR2 reporter assay (Figure 1A, Figure 2A-F).

Mutation of *lgt* has been associated with growth defects in cation-limiting conditions, human blood and mouse bronchoalveolar lavage fluid (14). Fewer planktonic BHN418 *lgt* mutant bacteria were also recovered from our NPE infection experiments (Figure 2C). Time course sampling of planktonic pneumococci grown with Detroit 562 cells revealed a minor growth defect for the BHN418 *lgt* mutant starting at 3 hpi but not for the TIGR4 *lgt* mutant (Figure 3A-B). To determine if the growth defect was dependent on the presence of NPE cells, time course sampling of planktonic BHN418 and its *lgt* mutant grown in infection medium and rich THY medium were performed. The growth defect was replicated in cell-free medium and is therefore not dependent on the presence of NPE cells (Figure 3C-D).

Our results indicate that inactivation of Lgt and therefore the lipoprotein processing pathway had greater consequences for BHN418 compared to TIGR4, except in their ability to trigger epithelial inflammation. These observations suggest that activation of the TLR2 pathway during pneumococcal-epithelial interactions is not dependent on the number of cell-associated or intracellular pneumococci. Additionally, within the timeframe of our assays, TLR2 signalling neither promotes nor inhibits epithelial microinvasion by *S. pneumoniae*.

**Figure 2.**
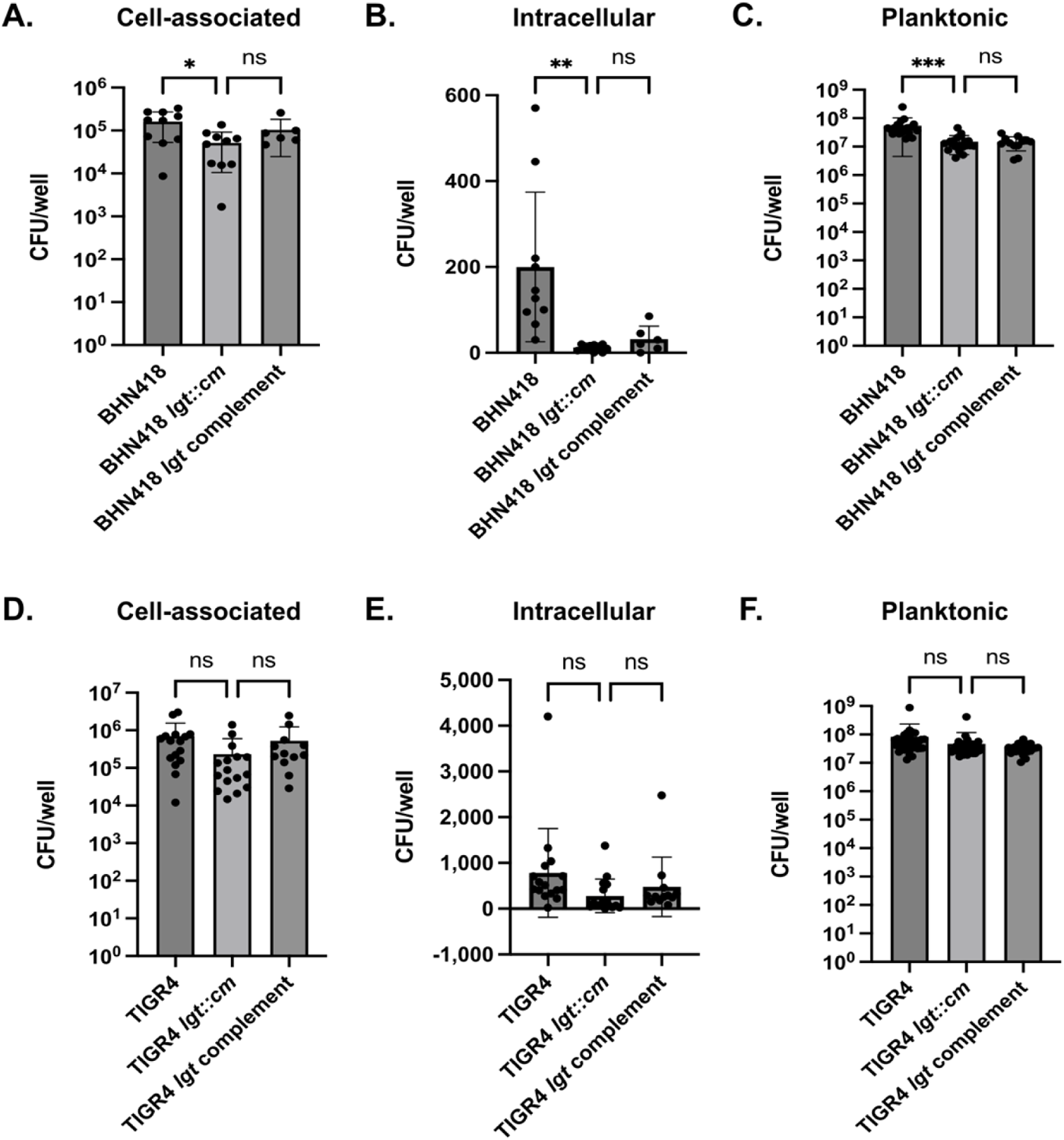
Mutation of *lgt* impaired epithelial microinvasion by *Streptococcus pneumoniae*, with greater defects for BHN418. (A-F) NPE microinvasion by WT and *lgt* mutants. Graphs show CFU numbers for BHN418-derived (A-C) and TIGR4-derived strains (D-F) associated with (A,D), internalised into (B,E), or growing in proximity with Detroit 562 NPE cells 3 hours post infection. * indicates *p* < 0.05, ** indicates *p* < 0.01, *** indicates *p* < 0.001.

**Figure 3.**
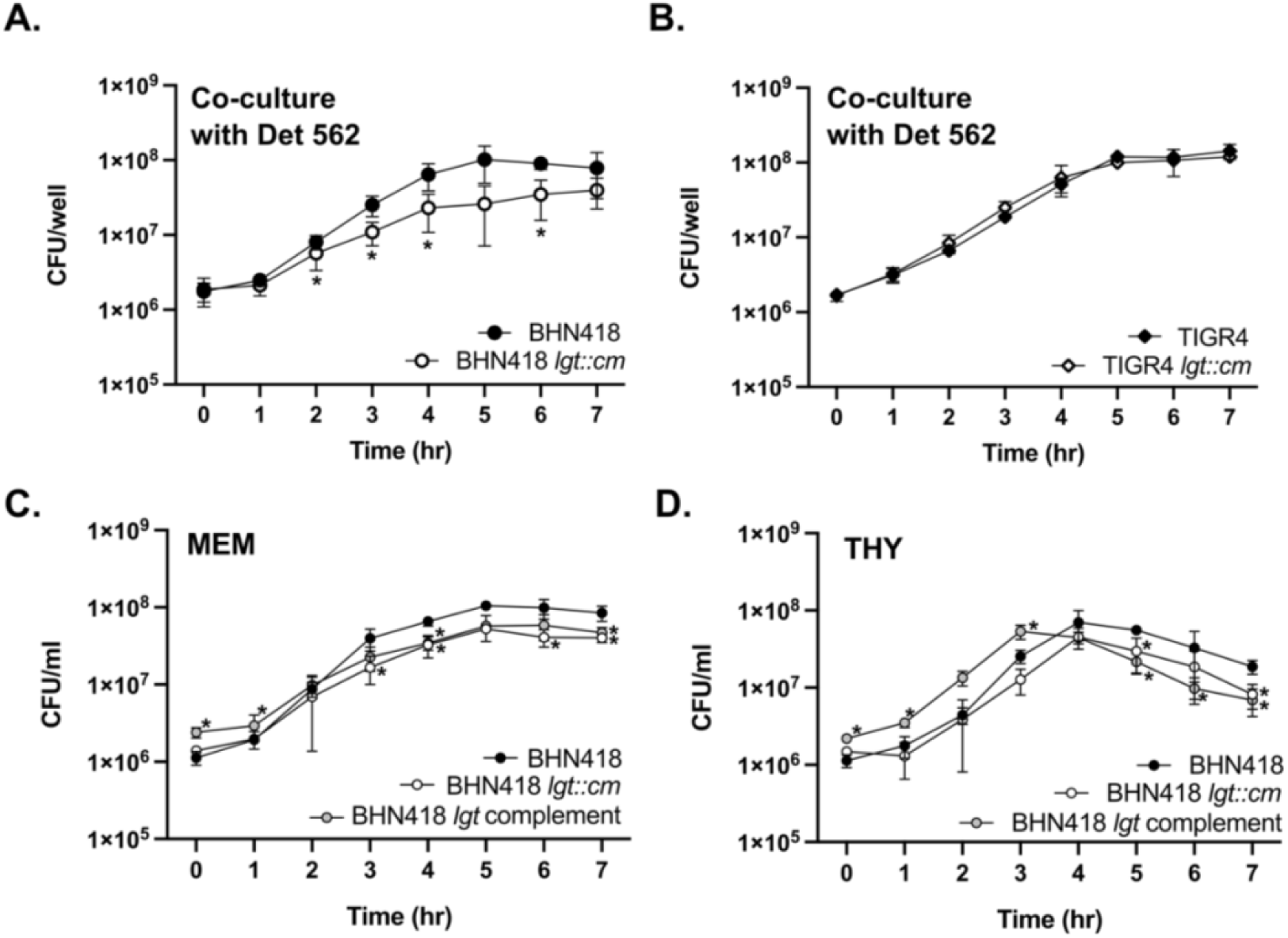
Growth of BHN418 *lgt::cm* compared to WT. (A-B) Growth of BHN418-derived (A) and TIGR4-derived (B) strains in infection medium (MEM + 1% FBS) in the presence of Detroit 562 NPE cells. (C-D) Growth of BHN418-derived strains in infection medium (C) and in the rich growth medium THY (D) in the absence of cells. * indicates *p* <0.05 by Student’s *t-*test.

### BHN418 encodes a novel lipoprotein absent in TIGR4

One potential explanation for the intraspecies differences in microinvasion upon *lgt* mutation is the presence of one or more lipoproteins in BHN418 which are absent in TIGR4. This lipoprotein may play a role as an adhesin and/or be important for nutrient transport and growth during infections. To address this hypothesis, we used a motif-based sequence toolkit, the MEME Suite, to compare the lipoprotein repertoires of BHN418 and TIGR4 (22). We identified 43 open reading frames (ORFs) in TIGR4 and 44 ORFs in BHN418 with gene products that fit the criteria for a lipoprotein (detailed in Materials and Methods) (Table 1). Of these putative lipoprotein ORFs, only one is present in BHN418 but not in TIGR4. There are no lipoprotein ORFs present in TIGR4 that are not also present in BHN418.

The lipoprotein encoded by BHN418 but not TIGR4, encoded by the gene with the locus tag RSS80_03595 and which we named <u>p</u>neumococcal <u>a</u>ccesssory lipoprotein <u>A</u> (PalA), comprises of 525 amino acids with sequence and structural homology to extracellular solute binding domain proteins that deliver substrates to ABC family transporters (Figure 4A-B). ABC transporters are multi subunit proteins comprising of two transmembrane permease domains and two cytoplasmic ATPase domains (23). Two genes encoding ABC transporter permease domain proteins, annotated as *yteP* and *araQ*, were found ∼3.3kb and ∼2.4kb upstream of *palA.* We were unable to locate ORF(s) encoding for the ATPase domain proteins in the 10kb region upstream or downstream of *palA*. Taken together, PalA likely binds to and delivers substrate(s) to YteP and/or AraQ. It is uncertain if YteP and/or AraQ co-opt the ATPase domains of ABC family transporters encoded elsewhere on the genome, in a similar strategy as the raffinose utilisation system, or no longer function as transporters (24, 25).

This genetic context suggests that *palA* is the fifth gene in an operon encoding for carbohydrate import and utilisation genes, which includes *yteP* and *araQ* (Figure 4A). The operon sits within an 11.5 kb region with ∼30% GC content, flanked by repetitive insertion sequences with homology to *IS630* elements. This region was likely acquired via horizontal gene transfer as the overall mean GC content of pneumococcal strains is around 40% (26). This putative genetic island is directly downstream of *spxB*, which encodes an important pneumococcal virulence factor involved in the production of H2O2 (27). Alignment of TIGR4 whole genome sequencing reads to the BHN418 genome revealed the absence of the entire putative island in the TIGR4 genome (Figure 4C) (26).

**Table 1.**
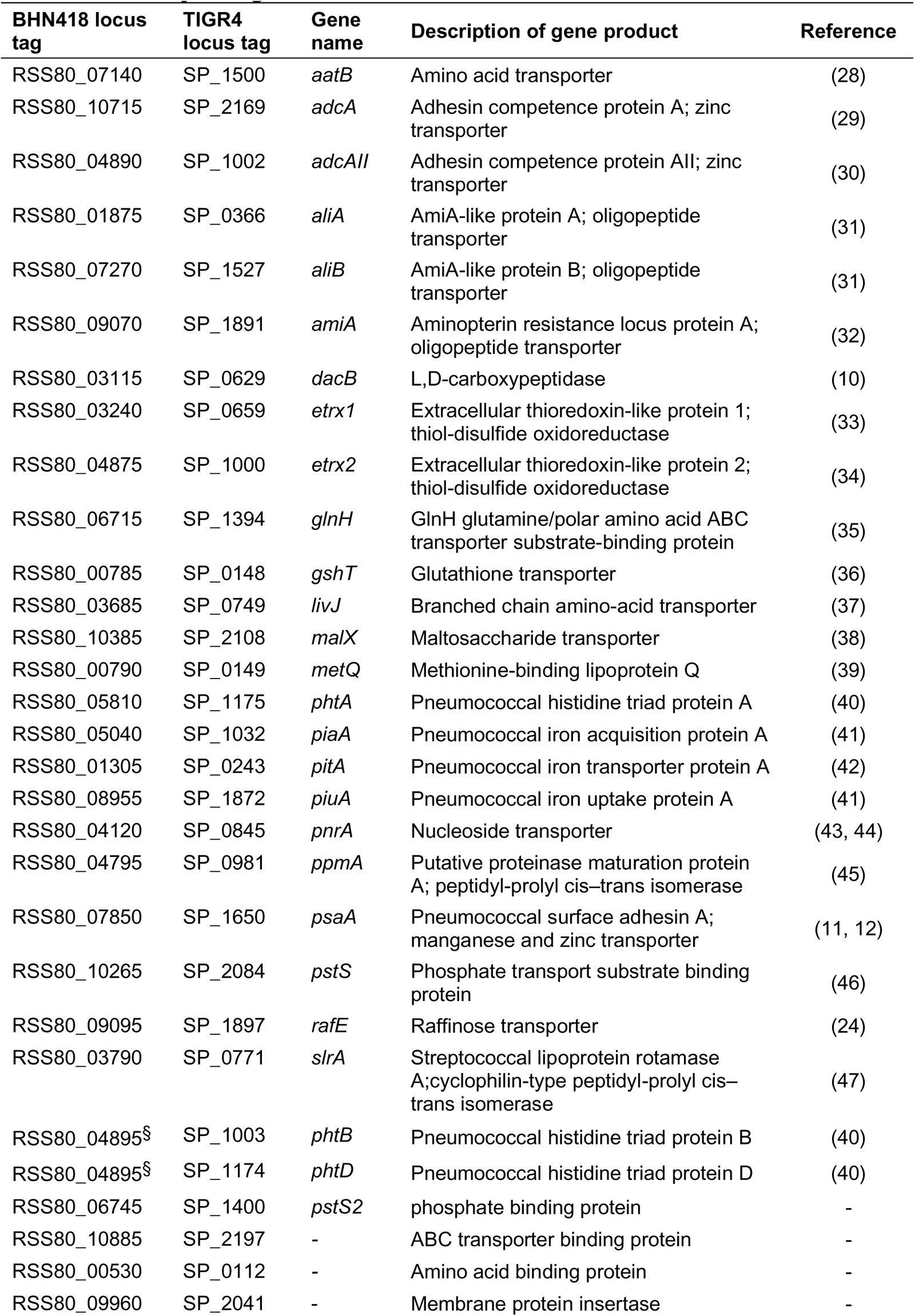
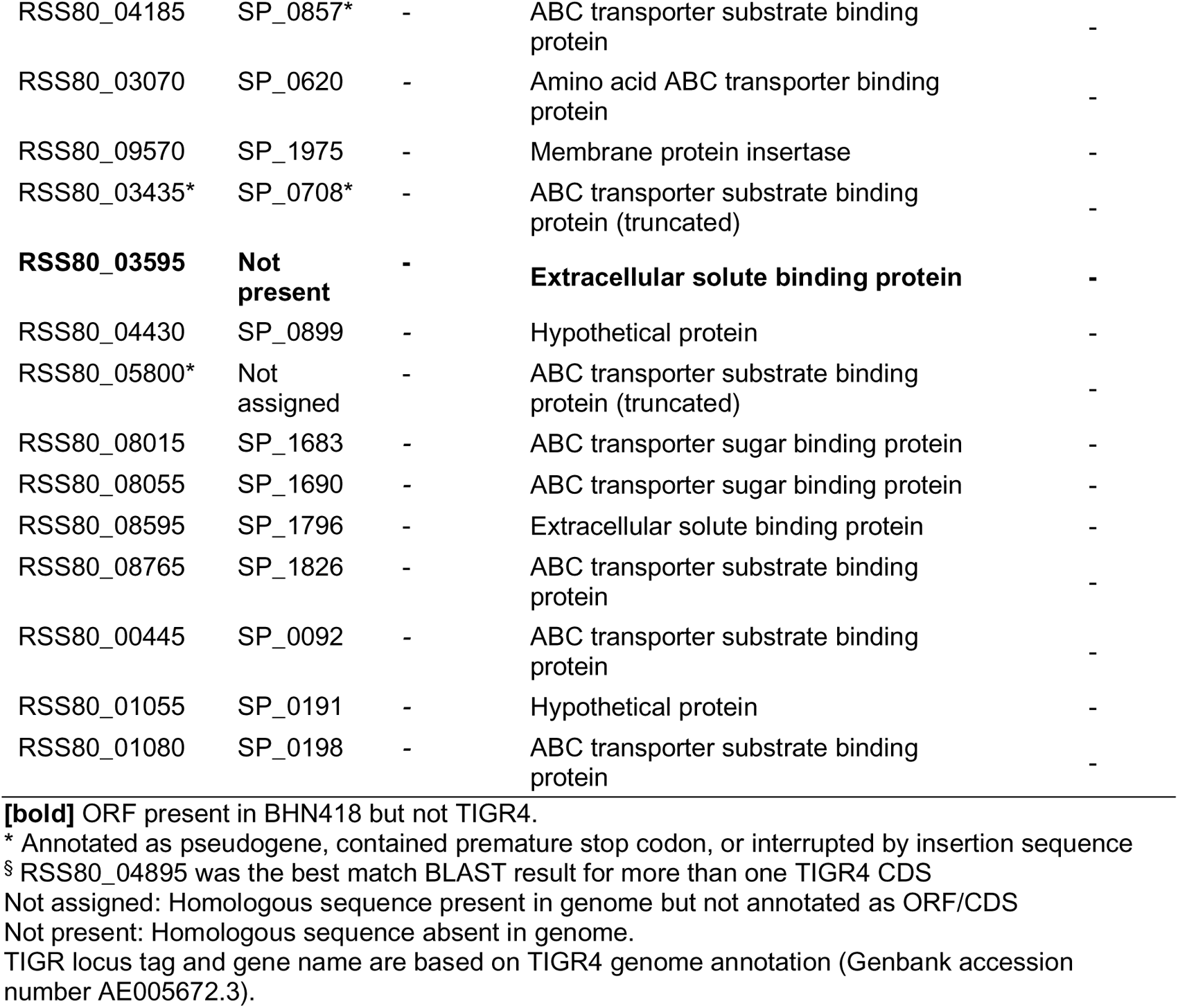
Lipoproteins encoded by TIGR4 and BHN418, identified bioinformatically using MEME suite.

**Figure 4.**
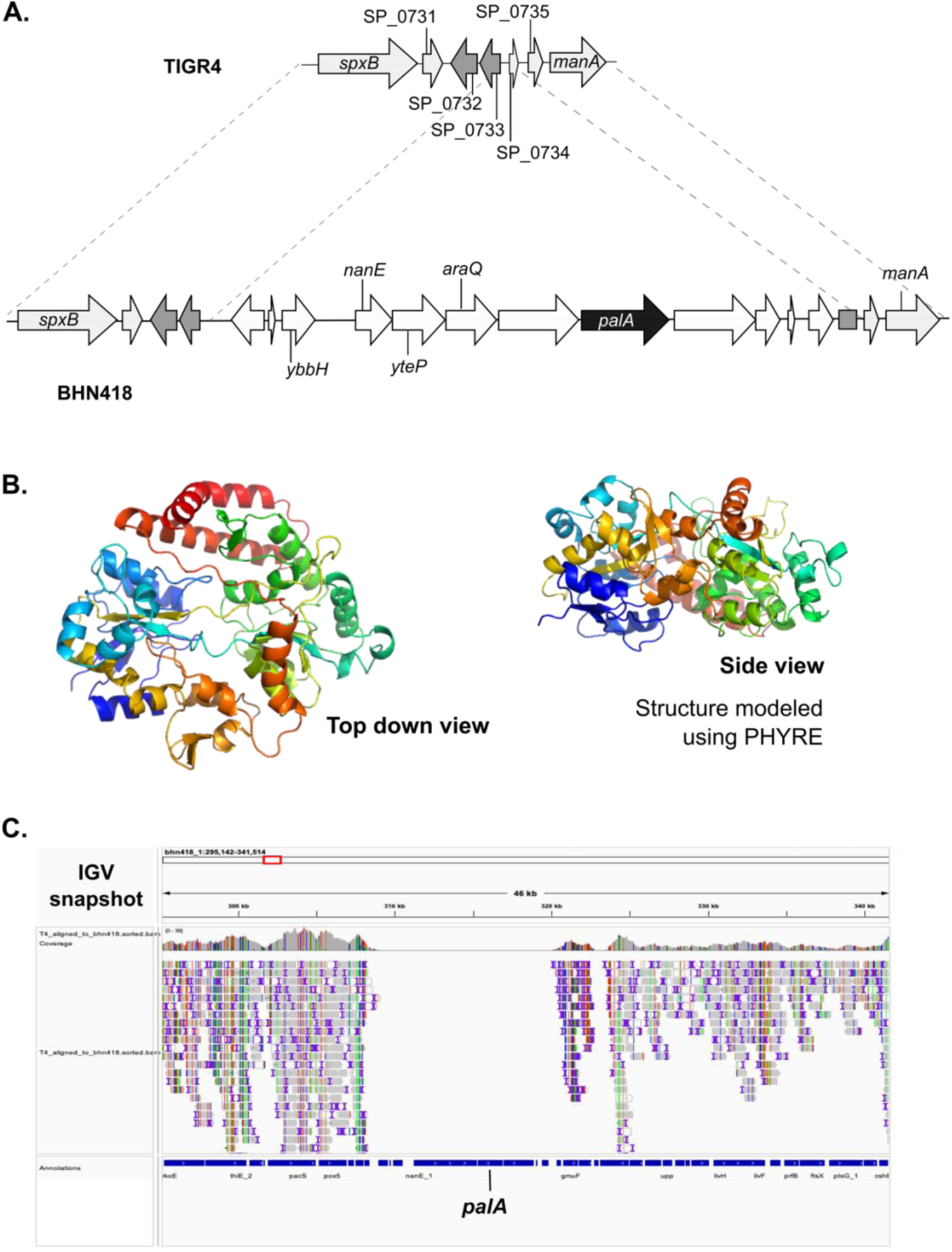
PalA is a lipoprotein present in BHN418 but not in TIGR4. (A) Genetic context of the *palA* gene (black arrow), which is within a multi-gene operon that is part of a putative genetic island. ORFs with homology to previously described genes are labelled with the gene name. (B) Predicted structure of PalA, modelled using PHYRE, which shows a barrel-like structure with a possible central ligand binding pocket. (C) IGV snapshot demonstrating lack of TIGR4 sequencing reads mapping to the putative *palA* genetic island.

To determine if *palA* is predominantly present in more carriage-type serotypes, such as serotype 6B, we examined the presence of *palA* in a well-curated dataset of 2806 carriage isolates from Malawi (48). 567 of these carriage isolates (20.5%) carry *palA* in their chromosome. Mapping the analysis results onto a hierarchal clustering (Newick) tree showed that *palA* is present in specific lineages, with no clear association to capsular serotypes or sequence types (genetic relatedness, visualised as neighbouring branches on a Newick tree) (Supplementary Figure 2) (48). However, presence of *palA* is enriched in certain serotypes, particularly serotype 6A (39/93, 41.9%), 6B (12/31, 38.7%), 10A (29/34, 85%), 15B (52.76, 68.4%), 16F (60/94, 63.8%), 23B (45/103, 43.7%), 35A (28/28, 100%) and 35B (60/114, 52.6%) (Supplementary Table 1). Additionally, the branching patterns of the phylogenetic tree for the Malawi carriage isolates supports the inference that *palA* and its associated genetic island were acquired via horizontal gene transfer and expanded in specific lineages (Supplementary Figure 2).

### PalA presence is enriched in carriage and ear isolates

Maintenance of this 11.5kb genetic island is potentially costly and suggests that the island confers some form of advantage to isolates that carry it. *S. pneumoniae* is capable of colonising and infecting multiple body sites including the nasopharynx, lungs, blood, CSF, meninges, and middle ear. We therefore examined 51,379 genomes in the BIGSdb database to determine if there is an association between the presence of *palA* and the isolation site of the strain (“source”) (49).

The *palA* gene presence is enriched in carriage isolates and in strains isolated from ear infections compared to strains isolated from IPD or lower respiratory tract disease (Table 2). More than half of serotype 22F and 6A strains isolated from the ear carried *palA* and approximately 28% of all serotype 22F and 6A genomes in the database carry *palA*, in contrast to the overall *palA* prevalence rate of 9.97% (Table 2). In the BIGSdb database, the only serotype 4 strain isolated from the ear carried *palA* in its genome. These observations suggest that *palA* and/or its putative genetic island may facilitate spread to and cause infection of the ear, although *palA*’s presence is not necessary for colonisation of the ear.

**Table 2.** Presence of *palA* in whole genome sequences of pneumococcal isolates on the BIGSdb database, stratified by site of isolation (“source”).

| Category | Source (BIGSdb label) | Proportion (%) |
| --- | --- | --- |
| Carriage | "nasopharynx", "pharynx", "sputum" | 14.57 (3114/21369) |
| Otitis | "ear swab", "middle ear fluid" | 13.72 (129/940) |
| Pneumonia | "lung aspirate", "sinus aspirate",<br>"bronchoalveolar lavage", "bronchi" | 8.96 (25/279) |
| Invasive | "blood", "cerebrospinal fluid", "joint<br>fluid", "pleural fluid" | 6.54 (1837/28075) |
| Eye/pus/others | "eye swab", "pus", "other" | 2.65 (19/716) |
| Overall |  | 9.97 (5124/51379) |

### Mutation of *palA* does not alter pneumococcal colonisation or microinvasion of the epithelium

To determine if PalA plays a role in epithelial microinvasion, we generated *palA* deletion and complementation mutants for testing in our NPE model. Although there is a small reduction in the number of planktonic bacteria, the numbers of epithelial-associated and intracellular BHN418 *palA::kan* were not significantly different to that of WT BHN418 (Figure 5A-C). Additionally, we did not observe a growth defect when BHN418 *palA::kan* was grown in THY or MEM (Figure 5D-E). Heterologous expression of *palA* in a serotype 23F strain naturally lacking the island (P1121) did not increase the microinvasion potential of the resulting strains and reduced the number of planktonic bacteria in the cell culture supernatant (Figure 6A-C). Moreover, the BHN418 *palA* knockout strains and the P1121 *palA* knock-in strains activated TLR2 signalling to similar levels as their respective wild-type strains (Figure 6D). We therefore conclude that presence of *palA* is not solely responsible for the observed strain-specific differences in Lgt-mediated epithelial microinvasion. Moreover, PalA does not contribute significantly to pneumococci’s ability to activate TLR2.

Mutation of *palA* attenuates nasopharyngeal colonisation density and duration in mice (14). We next asked if presence of *palA* confer a survival advantage in a more complex and immune-replete environment such as the murine nasopharynx. Outbred CD-1 female mice were intranasally inoculated with wild-type BHN418 and the *palA* mutant either singly or in a 1:1 competitive mix. After 7 days of colonisation, similar CFU numbers for WT BHN418 and the *palA::kan* were recovered from nasal washes (Figure 5F-G). Similar CFU numbers for BHN418 and *palA::kan* were also recovered in homogenised lungs and blood 24 hours post inoculation in a murine pneumonia model (Figure 5H-I). We conclude that presence of *palA* does not confer a colonisation advantage in the murine nasopharynx or in the progression to bacteraemic pneumonia.

**Figure 5.**
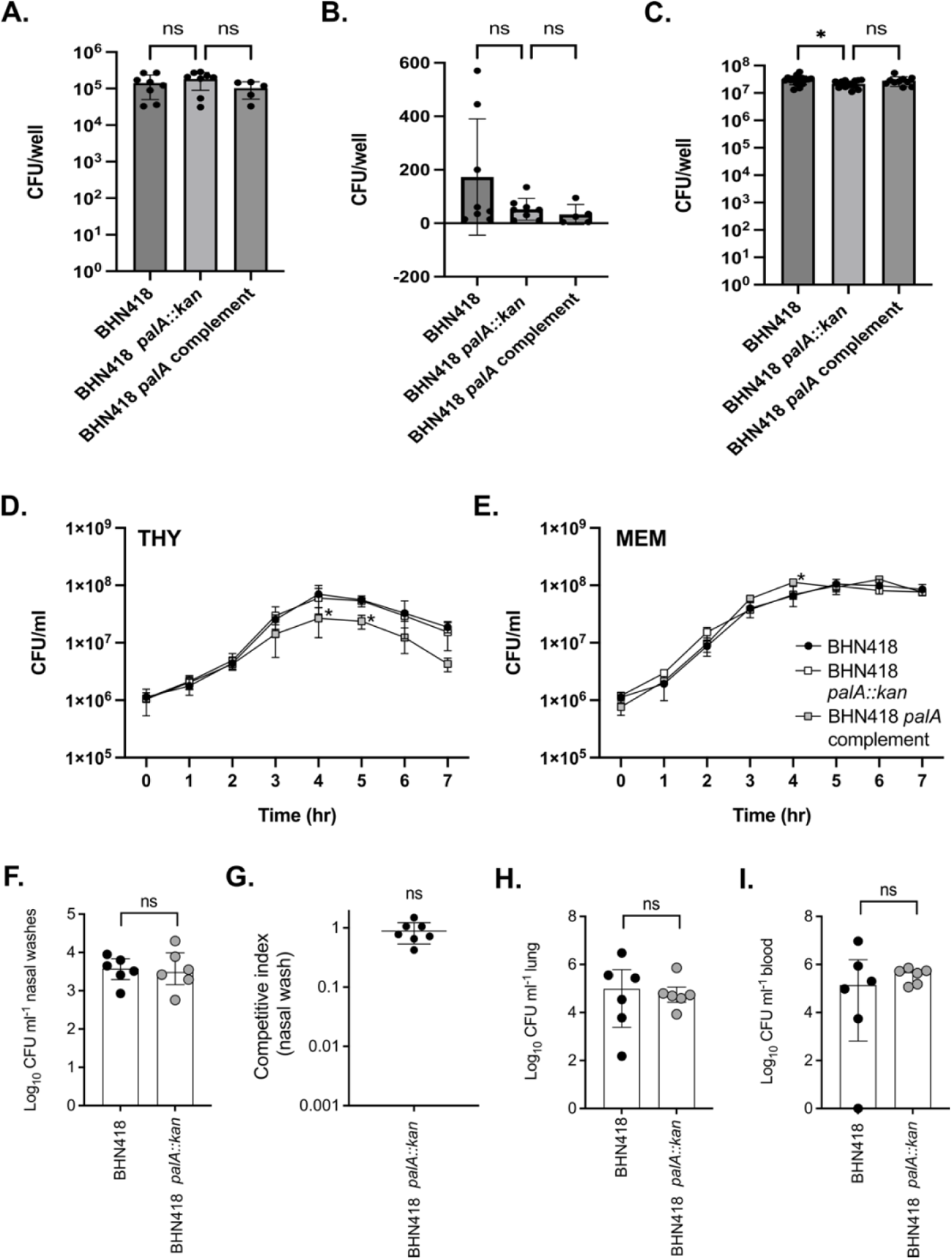
PalA was not essential for NPE microinvasion, murine colonisation and progression to disease. (A-C) NPE microinvasion by WT BHN418 and *palA* mutants, measured as NPE-associated bacteria (A), internalised bacteria (B) and planktonic bacteria growing in proximity with Detroit 562 NPE cells 3 hours post infection. (D-E) Growth of WT BHN418, the *palA* knock out and complementation mutants in THY (D) and infection medium (E). (F-I) Recovery of pneumococci from mice intranasally inoculated with WT BHN418 and *palA::kan* mutant, recovered from nasal washes when inoculated singly (F) or competitively in a 1:1 ratio (G), as well as from the lungs (H) and bloodstream (I) when tested on a pneumonia model. * indicates *p* < 0.05.

**Figure 6.**
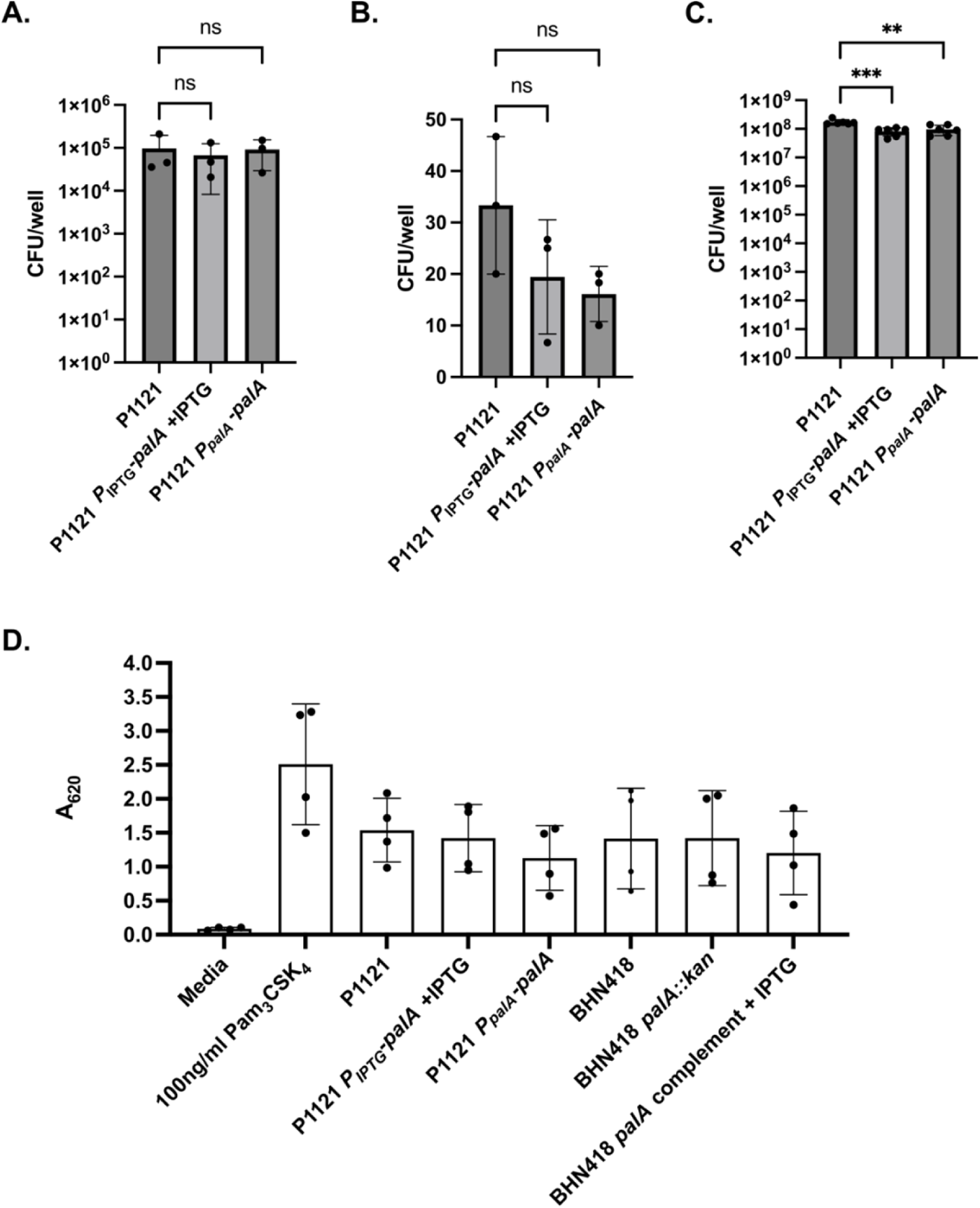
Heterologous expression of *palA* in P1121 (serotype 23F) did not increase epithelial microinvasion or TLR2 signalling. (A-C) NPE microinvasion by WT P1121 and *palA* expression mutants, measured as NPE-associated bacteria (A), internalised bacteria (B) and planktonic bacteria growing in proximity with Detroit 562 NPE cells 3 hours post infection. (D) SEAP reporter readout from HEK-Blue^TM^ hTLR2 reporter cells treated with pneumococcal strains at MOI 10 for 16 hours.

## DISCUSSION

In this study, we demonstrated that pneumococcal lipoproteins trigger inflammation during epithelial colonisation at least partially via the TLR2-dependent pathway. We have previously shown that epithelial microinvasion occurs in the absence of disease and that there is heightened epithelial inflammation around the time of pneumococcal clearance in controlled human infection (2). In murine models, mutation of *lgt* reduces carriage duration and attenuates disease, associated with a concomitant reduction in inflammatory and immune responses (8, 9, 50). However, although we have shown that BHN418 *lgt::cm* but not TIGR4 *lgt::cm* was significantly attenuated in epithelial adherence and microinvasion compared to their respective wild-type strains, this does not appear to be TLR2-dependent or due to differential lipoprotein repertoires encoded by the strains.

We additionally observed that presence of *lgt* and therefore TLR2 activation heightens the epithelial interferon response elicited by pneumococcal microinvasion in the absence of immune cells. Induction of the interferon pathway upon pneumococcal challenge is thought to be dependent on sensing of intracellular pneumococci, pneumococcal DNA or cellular DNA damage by the infected cells (51–56). In infant mice co-infected with pneumococci and influenzae, interferon signalling increases bacterial shedding while protecting against invasive disease, while TLR2 signalling limits bacterial shedding and transmission (18, 57, 58). Since TLR2 signalling augments the interferon response by human nasopharyngeal epithelial cells during mono-pneumococcal infection, it is unclear how these two pathways mediate the outcomes of pneumococcal microinvasion, colonisation or progression to disease in people. Nonetheless, our results suggest that mucosal innate immunity could be targeted alongside the induction of an adaptive immune response to prevent pneumococcal colonisation, transmission and invasive disease.

We have implicated lipoprotein expression in intraspecies differences in NPE cell microinvasion but our genetic and mutational analysis lead us to suggest that this is not due to differences in lipoprotein repertoire, but rather due to differential post-translational lipidation. The lipoprotein PsaA functions as a bacterial adhesin which binds to host cell E-cadherin (59). In theory, we should have observed a significant reduction in the number of epithelial-associated bacteria for both the TIGR4 and BHN418 *lgt* mutants compared to wild-type due to the loss of PsaA surface presentation; however, our data did not reflect this hypothesis. Similar subtle effects have been seen for example when mutating the lipoprotein encoding gene *dacB* but not *lgt* alters murein sacculus composition, and when mutating *lgt* results in strain-dependent variable effects on growth in rich medium (5, 9, 14, 50, 60). The observation that genetic complementation of *lgt* in BHN418 restores pneumococcal ability to trigger epithelial inflammation but not WT levels of microinvasion further suggests that regulation of lipoprotein processing may be more complex than previously thought. The assumption that Lgt and other lipoprotein modification proteins are constitutively expressed and active, and all 30+ lipoproteins are processed at similar rates may not be entirely correct. Proteomics and immunoblotting analyses showed that abundance of specific lipoproteins may increase, decrease or show no change when lipoprotein processing is disrupted in *S. pneumoniae*, although no clear patterns were apparent (5, 13). We therefore speculate that unlike Gram-negative bacteria, for which lipoprotein processing is essential, pneumococci and other Gram-positive bacteria compensate for the loss of Lgt by differentially regulating expression of lipoprotein encoding genes, which in turn is regulated by external stimuli in nasopharyngeal niche (61, 62).

While investigating the intraspecies variation in the role of *lgt* in microinvasion, we discovered a previously uncharacterised lipoprotein encoding gene (*palA*) and its associated genetic island. Although *palA* presence is enriched in carriage and otitis media isolates, we have not been able to demonstrate a clear role for PalA in epithelial microinvasion, or in a murine model, colonisation or disease. PalA is predicted by sequence and structure homology to be involved in carbohydrate or sugar transport, although we have yet to identify a substrate for PalA. Raffinose metabolism has been shown to contribute to lung versus ear tropism in serotype 3 and serotype 14 strains (25). It is therefore possible that *palA* functions in promoting niche specialisation by facilitating uptake and metabolism of uncommon sugars, such as raffinose, found in the nasopharynx or middle ear.

In conclusion, we demonstrated a role for pneumococcal surface lipoproteins in triggering epithelial inflammation and augmenting interferon signalling in response to pneumococcal-epithelial interactions. We show that pneumococcal lipoproteins mediate microinvasion in a strain-dependent manner, which may explain the significant attenuation in carriage duration and disease with *lgt* mutants reported by others (8, 14, 50). Additionally, we have characterised a novel accessory lipoprotein likely acquired through horizontal gene transfer but rejected the hypothesis that this lipoprotein contribute to strain differences in pneumococcal epithelial microinvasion. Instead, we postulate that differential regulation of lipoprotein gene expression responding to the nasopharyngeal niche regulate this microinvasion process.

## MATERIALS AND METHODS

### Bacterial growth and maintenance

*Streptococcus pneumoniae* strains were grown on Columbia agar base with 5% defibrinated horse blood (CBA plates; EO Labs, Oxoid) or statically in Todd-Hewitt broth supplemented with 0.5% yeast extract (THY; Oxoid) at 37°C, 5% CO_2_. Where appropriate, growth medium was supplemented with antibiotics at the following concentrations: chloramphenicol (10 µg/ml), erythromycin (0.5 µg/ml), kanamycin (250 µg/ml). Working stocks for infections were prepared by freezing THY cultures at OD_600_ 0.3-0.4 with 10% glycerol. NEB® Stable competent *Escherichia coli* derived strains were grown in LB broth or LB agar (Difco) supplemented with ampicillin (200 µg/ml) where appropriate. Bacterial strains used in this paper are listed in Table 3.

### Bacterial genetic manipulation

*S. pneumoniae* were genetically manipulated using a competence stimulating peptide (CSP)-mediated transformation assay (63). Briefly, pneumococci were grown in THY pH 6.8 supplemented with 1 mM CaCl_2_ and 0.02% BSA at 37°C, 5% CO_2_ to OD_600_ 0.01-0.03, pelleted and resuspended in 1/12 volume THY pH 8.0 supplemented with 1 mM CaCl_2_ and 0.2% BSA. A total of 400 ng CSP (Cambridge Biosciences; CSP-2 for TIGR4; 1:1 ratio of CSP-1:CSP-2 for BHN418) was added to the bacterial suspension and incubated at RT for 5 mins. The suspensions were then mixed with ∼300 ng transforming DNA, incubated at 37°C, 5% CO_2_ for two hours and plated on CBA plates supplemented with relevant antibiotics. Antibiotic resistant transformants were screened using colony PCR and confirmed by sequencing.

Transforming DNA for generating *lgt::cm* and *palA::kan* mutants were generated using overlap-extension PCR. Complementation and expression constructs were generated by inserting the target gene into the complementation plasmid pASR103 or pPEPY (64), which allows for integration of the construct at a chromosomal ectopic site. Plasmids used are listed in Table 3, while primers are listed in Supplementary Table 2.

### Cell culture

Detroit 562 (ATCC® CCL-138^TM^ human pharyngeal carcinoma epithelial cells) were expanded and maintained in MEMα (Gibco^TM^ 22561021) supplemented with 10% heat-inactivated FBS (HI-FBS; LabTech FB-1001/500 or Gibco 10438-026) at 37°C, 5% CO_2_. HEK-Blue^TM^ hTLR2 reporter cells (Invivogen, hkb-htlr2) were expanded and maintained in DMEM (4.5 g/L glucose, 2mM glutamine, sodium pyruvate) supplemented with 10% HI-FBS at 37°C, 5% CO_2_. Per manufacturer’s instructions, DMEM growth medium was supplemented with 100 µg/ml normocin^TM^ and/or 1X HEK-Blue^TM^ Selection (Invivogen) where appropriate.

### NPE infections

Adherence-invasion infections of confluent Detroit 562 cells with *S. pneumoniae* strains were performed at MOI 20 (P1121/23F derived strains) or MOI 10 (all others) for 3 hours. Working bacterial stocks were thawed, centrifuged to remove freezing medium and resuspended in infection medium (MEMα with 1% HI-FBS) to the appropriate CFU. 1 ml bacterial suspension were added to each well containing confluent Detroit 562 cells. Plates were incubated statically at 37°C, 5% CO_2_ for 3 hours, after which 10 µl of the supernatant were removed for CFU enumeration. For adherence assays, cells were washed thrice with PBS, lysed with cold 1% saponin (10 min incubation at 37°C, followed by vigorous pipetting), and 10 µl cell lysate removed for CFU enumeration. For invasion assays, cells were washed thrice with PBS, incubated with 0.5 ml infection medium supplemented with 200 µg/ml gentamicin at 37°C, 5% CO_2_ for 1 hour to kill extracellular bacteria, followed by 3x PBS wash, lysis with 1% saponin and CFU enumeration. Experiments were performed at least thrice on different days (n≥3 biological replicates) with technical duplicates. Statistical significance was determined using one-way ANOVA with Bonferroni’s multiple comparison test.

To harvest RNA for qPCR, confluent Detroit 562 cells were treated with synthetic agonists or infected with *S. pneumoniae* strains at MOI 10 for 6 hours. Briefly, working bacterial stocks were thawed, centrifuged to remove freezing medium and resuspended in infection medium (MEMα with 1% HI-FBS) to the appropriate CFU. Bacterial suspensions, infection medium (negative control), or infection medium supplemented with synthetic agonists (20 µg/ml Poly(I:C)) (TLR3 agonist, Bio-Techne) were added to each flask. Flasks were incubated statically at 37°C, 5% CO_2_ for 6 hours, after which 10 µl were removed for CFU enumeration. Detroit 562 cells were washed thrice with PBS and harvested by scraping into 300 µl RNA*later* (ThermoFisher). For each treatment condition, RNA harvesting was performed at least thrice on different days (n≥3 biological replicates) without technical replicates.

For growth curve experiments, *S. pneumoniae* strains were seeded into 1 ml THY or 1 ml infection medium (MEMα with 1% HI-FBS, LabTech) with and without confluent Detroit 562 cells in 12-well plates at a similar CFU number as used in infection experiments. Plates were incubated at 37°C, 5% CO_2_ for 7 hours, with aliquots taken for CFU enumeration every hour. CFU growth curves were performed at least thrice on different days (n≥3 biological replicates) without technical replicates. Statistical significance was determined using Student’s *t-*test assuming equal variance.

### qPCR

RNA from epithelial cells stored in RNA*later* were extracted using RNeasy Mini kit (Qiagen) according to manufacturer instructions. Carryover DNA was removed with TURBO DNA-*free* kit (Ambion), and cDNA generated using LunaScript® RT Supermix kit (NEB). qPCR was performed using Luna® Universal qPCR Master Mix (NEB) in technical triplicates with primers specific for *GAPDH*, *CXCL10*, *IFNB1*, *IFNL1,* and *IFNL3* (Supplementary Table 2). Whenever possible qPCR primers were designed to span exon-exon junctions. Cycling conditions are as follows: 95°C for 5 mins, 40 cycles of 95°C for 15 secs and 60.5°C for 45 secs, with a plate read at the end of each cycle. Data was analysed using the 2^ΔΔCt^ method, with media only control and *GAPDH* levels for normalization. Statistical significance was determined using Student’s *t-*test assuming equal variance.

### HEK-Blue hTLR2 reporter assay

HEK-Blue^TM^ hTLR2 secreted alkaline phosphatase (SEAP) reporter assays were performed according to manufacturer instructions (Invivogen, hkb-htlr2). Briefly, HEK-Blue^TM^ hTLR2 cells, *S. pneumoniae* and control reagents were resuspended or diluted in pre-warmed HEK-Blue^TM^ Detection medium (Invivogen). 5 x 10^4^ HEK-Blue^TM^ hTLR2 cells were mixed with 5 x 10^5^ CFU *S. pneumoniae* (MOI 10) and incubated for 16 hours at 37°C, 5% CO_2_. SEAP activity was then measured spectroscopically at A_620_. 100 ng/ml of Pam_2_CSK_4_ and Pam_3_CSK_4_ (TLR2 agonist, Bio-Techne) were used as positive controls, while bacterial-free medium was used as negative control. Experiments were performed at least thrice on different days (n≥3 biological replicates) with technical triplicates. Statistical significance was determined using one-way ANOVA with Bonferroni’s multiple comparison test.

### Lipoprotein prediction using MEME suite

Amino acid sequences of thirty-nine published D39 lipoproteins were used with the motif discovery tool MEME to identify pneumococcal lipoprotein motif(s) (5, 13, 22). The top two MEME results were combined to obtain motif: L[LA][AS][AL]LXL[AV]A**C**[SG][NQS], a modified extension of the minimal lipobox motif LAG**C** (5).

The obtained motif was used with the motif scanning tool FIMO to identify lipoproteins in the genomes of *S. pneumoniae* TIGR4, BHN418 and D39, with the latter used for quality control (65). Match *p*-value was set to 0.001. FIMO results were further filtered with the following criteria: (i) presence of the lipidated cysteine residue in the motif, (ii) presence of motif in the first 70 a.a. of the sequence, iii) positive prediction as lipoprotein by SignalP-6.0 (66).

### Genomic analysis

Presence of *palA* and its associated genetic island were determined using Local-BLAST (BLASTN, TBLASTN) for the Malawian carriage dataset (n=51,379) and serotype 23F strain P1121 (67). The built-in BLAST tool on pubmlst.org was used for analysis of the BIGSdb dataset (49). BLASTN and TBLASTN tools on the NCBI database were used to identify *palA* and PalA homologues in non-pneumococcal species (67, 68). BLAST results were exported in csv format and further analysed using R (v3.6.0) in RStudio (http://www.rstudio.com/). Presence/absence of *palA* was annotated onto a Newick tree showing phylogeny of the Malawian carriage strains by metabolic type and visualized using iTOL (48, 69). Potential gene functions were inferred through the results of BLASTP and NCBI Conserved Domain Database searches (67, 68, 70).

The BHN418 genome assembly was generated by combining long read sequencing (PacBio) and short read sequencing (Illumina) methods which resulted in a single contiguous chromosome of BHN418 of length 2,107,426 bp. *De novo* assembly was performing using the Unicycler v0.4.8 pipeline in bold mode, quality assessed using QUAST v5.1.0rc1 and annotated using Bakta v1.8.2 as described previously (71–74). TIGR4 sequencing reads were aligned to the BHN418 genome using Samtools v1.14 and visualised using IGV v2.16.1 (26, 68).

### TLR2 transcriptional module analysis

TLR2-mediated transcriptional activity in Detroit 562 cells infected with TIGR4 and BHN418 for 3 hours were determined using published RNAseq data (2). We generated a transcriptional module reflective of TLR2 activity derived from genes overexpressed in fibroblasts stimulated with TLR2 agonists Pam_2_CSK_4_ and/or FSL-1 for 6 hours relative to unstimulated controls (>1.5 fold; paired *t*-test with α of p<0.05 without multiple testing correction) (Gene Expression Omnibus (GEO) dataset GSE92466) (Supplementary Table 1) (19). Module expression was determined by calculating the geometric mean expression of all constituent genes found in the analysed RNAseq dataset. Performance was validated using data derived from Acute Myeloid Leukemia cells (GEO datasets GSE92744) and CD14+ monocytes stimulated with Pam3CSK4 (GEO dataset GSE78699) (Supplementary Figure 1) (75, 76).

### Murine experiments

Outbred female CD1 mice (Charles River Laboratories) were inoculated intranasally under anaesthetic (isoflurane) with 1 x 10^7^ CFU bacteria (n=6 for single inoculation colonisation and pneumonia model, n=7 for competition experiment). For colonisation experiments, nasal washes were performed 7 days post infection using 1 ml PBS. For pneumonia model, mice were sacrificed 24 hpi and bacteria recovered from the blood and homogenized lungs. CFU numbers were enumerated using CBA supplemented with 4 µg/ml gentamicin, with additional 250 µg/ml kanamycin where appropriate. All animal procedures were approved by the local ethical review process and conducted in accordance with the relevant UK Home Office approved project license (PPL70/6510). Mice were housed for at least one week under standard conditions before use. Randomisation or blinding was not performed for these experiments. Statistical significance was determined using Mann-Whitney test.

### Data availability

BHN418 genome was deposited to NCBI with accession number PRJNA1022026. TIGR4 sequencing reads were downloaded from NCBI Sequence Reads Archive (accession SRX6259281), while P1121 reads were downloaded from the EMBL-EBI database (accession ERS1072059) (77, 78). D39 and TIGR4 whole genome assemblies were downloaded from NCBI GenBank database (accession numbers CP000410.2 and AE005672.3, respectively) (79). All other genomic sequences used were hosted on the PubMLST Pneumococcal Genome Library (https://pubmlst.org/organisms/streptococcus-pneumoniae/pgl) (48, 49). RNAseq data used in the TLR2 transcriptional module expression analysis were obtained from the ArrayExpress database (accession E-MTAB-7841) (6).

**Table 3.** Bacterial strains and plasmids used in this study.

| Designation | Genotype/ Description | Source |
| --- | --- | --- |
| <u>Strains</u> |  |  |
| TIGR4 | WT Serotype 4 isolate | (26) |
| BHN418 | WT Serotype 6B isolate | (16) |
| P1121 | WT Serotype 23F isolate | (2) |
| ECSPN100 | TIGR4 *lgt::cm* | This work |
| ECSPN106 | TIGR4 *lgt::cm P<sub>IPTG</sub>-lgt-erm* (TIGR4 *lgt* complementation) | This work |
| ECSPN200 | BHN418 *lgt::cm* | This work |
| ECSPN210 | BHN418] *lgt::cm P<sub>IPTG</sub>-lgt-erm* (BHN418 *lgt* complementation) | This work |
| ECSPN211 | BHN418 *palA::kan* | This work |
| ECSPN213 | BHN418 *palA::kan P<sub>IPTG</sub>-palA-erm* (BHN418 *palA* complementation) | This work |
| ECSPN400 | P1121 *P<sub>IPTG</sub>-palA-erm* | This work |
| ECSPN401 | P1121 *P<sub>palA</sub>-palA-kan* | This work |
| <u>Plasmids</u> |  |  |
| pASR103 | Complementation construct with an IPTG inducible promoter and *erm* selectable marker | (64) |
| pPEPY | Complementation construct with a *kan* selectable marker | (64) |
| pEMcat | Minitransposon plasmid; source of *cm<sup>R</sup>* cassette | (80) |
| pABG5 | Cloning plasmid; source of *kan<sup>R</sup>* cassette | (81) |
| pEC210 | TIGR4 *lgt* coding region cloned into pASR103 | This work |
| pEC211 | BHN418 *lgt* coding region cloned into pASR103 | This work |
| pEC213 | BHN418 *palA* coding region cloned into pASR103 | This work |
| pEC213 | BHN418 *palA* promoter and coding region cloned into pPEPY | This work |

## ACKNOWLEDGEMENTS

This study was funded by a Medical Research Council grant (MR/T016329/1) awarded to RSH and JSB which supported JMC and CMW. JMC and MB were also supported by funding from NIHR Global Health Research Unit on Mucosal Pathogens at UCL, commissioned by the National Institute for Health Research using Official Development Assistance (ODA) funding. GP is supported by funding from the UCLH NIHR Biomedical Research Centre. RSH is a NIHR Senior Investigator. The views expressed are those of the authors and not necessarily those of the NIHR.

