## Supplemental materials for "Bacterial surface lipoproteins mediate epithelial microinvasion by *Streptococcus pneumoniae*"

**A.**

| Genes in TLR2 module |
|---|
| MIR21 |
| AQP9 |
| NFKBIZ |
| CCL7 |
| GBP1 |
| IRAK2 |
| METTL7A |
| PLA2G4A |
| EDNRA |
| TMEM140 |
| IFIH1 |
| PTGES |
| IER3 |
| OAS3 |
| SLC39A14 |
| CXCL5 |
| HERC6 |
| NFKBIA |
| BTN3A3 |
| SOD2 |
| IL8 |
| CXCL1 |
| CXCL6 |
| MIR302C |
| TNFAIP2 |
| MAP3K8 |
| GBP2 |
| RELB |
| LRRN3 |
| C8orf4 |
| CCL2 |
| LOC100134000 |
| DTX3L |

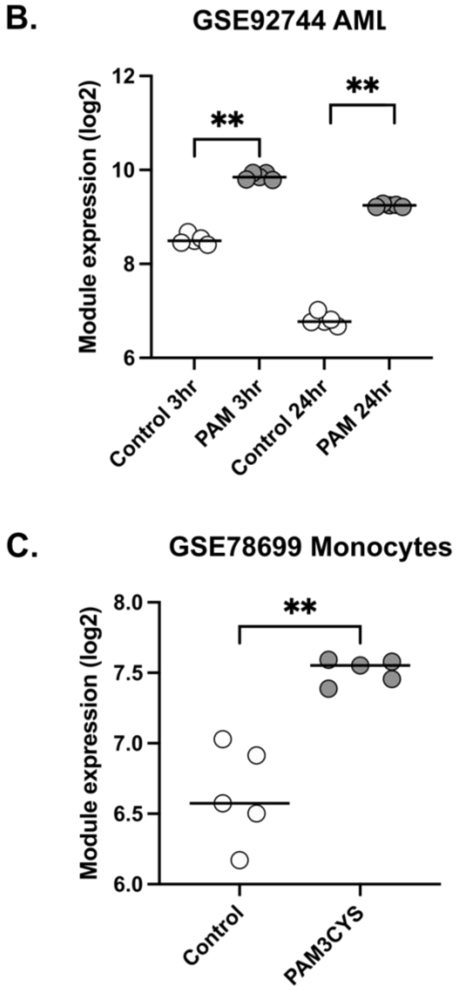


**Supplementary Figure 1. TLR2 transcriptional module.** Module was derived from genes overexpressed in fibroblasts stimulated with TLR2 agonists Pam$_2$CSK$_4$ and/or FSL-1 for 6 hours relative to unstimulated controls, listed in (A). Performance was validated using RNAseq data derived from Acute Myeloid Leukemia cells (B) and monocytes stimulated with Pam$_3$CSK$_4$. Statistical testing was performed using Mann-Whitney test. ** $p<0.01$.

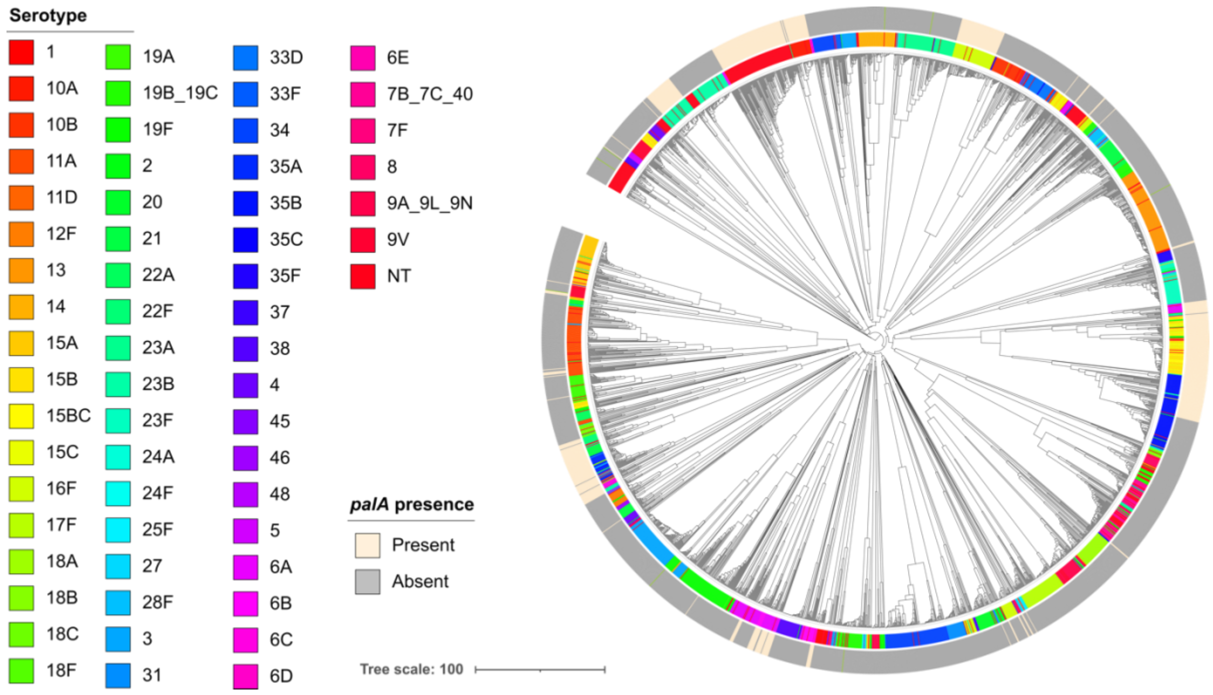


**Supplementary Figure 2. *palA* is encoded by diverse pneumococcal strains, with no clear association to specific serotypes.** Presence of *palA* mapped onto a hierarchal clustering tree based on metabolic type, constructed using whole genome sequences of 2806 carriage isolates from Malawi (48).

**Supplementary Table 1. Presence of *palA* in the whole genome sequences of 2806 carriage isolates from Malawi, stratified by serotype.**

| Vaccine type | Serotype | *palA* prevalence (%)<br>(*palA* present/total isolates) |
|---|---|---|
| *Overall dataset* | | 20.21 (567/2806) |
| Vaccine type | *All VT* | 11.11 (77/693) |
| (VT) | **6A** | **41.94 (39/93)** |
| | **6B** | **38.71 (12/31)** |
| | 9V | 14.29 (6/42) |
| | 19F | 5.8 (8/138) |
| | 18C | 4.76 (1/21) |
| | 3 | 4.44 (6/135) |
| | 19A | 4.35 (2/46) |
| | 23F | 4.11 (3/73) |
| Non-vaccine type | *All NVT* | 20.70 (362/1749) |
| (NVT) | **35A** | **100 (28/28)** |
| | **22F** | **85.71 (12/14)** |
| | **10A** | **85.29 (29/34)** |
| | **15BC** | **71.43 (15/21)** |
| | **15B** | **68.42 (52/76)** |
| | **16F** | **63.83 (60/94)** |
| | **35B** | **52.63 (60/114)** |
| | **15C** | **48.15 (13/27)** |
| | **23B** | **43.69 (45/103)** |
| | **21** | **20.29 (14/69)** |
| | 8 | 10 (2/20) |
| | 11A | 9.43 (10/106) |
| | 15A | 6.02 (5/83) |
| | 28F | 4.35 (1/23) |
| | 18A | 3.77 (2/53) |
| | 20 | 2.13 (1/47) |
| | 9A_9L_9N | 2.08 (1/48) |
| | 34 | 0.77 (1/130) |
| Nontypable (NT) | **NT** | **35.16 (128/364)** |

Serotypes shown are represented by ≥10 clinical isolates with at least one isolate carrying *palA*. Serotypes in **bold** have above average *palA* carriage rate.

**Supplementary Table 2. Primers used in this study. Restriction enzyme sites are underlined.**

| Primer Designation | Sequence (5'-3') | References |
|---|---|---|
| *Used in qPCR* | | |
| gapdh-RT-F | CGGATTTGGTCGTATTGG | This work |
| gapdh-RT-R | AGATGGTGATGGGATTTC | This work |
| CXCL10-RT-F | CCTGCTTCAAATATTTCCC | This work |
| CXCL10-RT-R | CCTTCCTGTATGTGTTTGGA | This work |
| ifnb-RT-F | CTTGGATTCCTACAAAGAAGC | This work |
| ifnb-RT-R | CATCTCATAGATGGTCAATGC | This work |
| IFNL1 RT-PCR F | GCCTCCTCACGCGAGACCTC | (82) |
| IFNL1 RT-PCR R | GGAGTAGGGCTCAGCGCATA | (82) |
| IFNL3 RT-PCR F | TGGCCCTGACGCTGAAGGTT | (82) |
| IFNL3 RT-PCR R | CGTGGGCTGAGGCTGGATAC | (82) |
| *Used to generate transforming DNA for making ECSPN100, ECSPN200.* | | |
| Sp1411F | GAGTCATCAAGAGCTTCGG | (14) |
| Cm-1411R | GCCTAATGACTGGCTTTTATAAATGTTAGAAGTTGCATATATTC | (14) |
| Cm-1413F | ACATTATCCATTAAAAATCAAATCAAGCATTTTGCACCTCATTT | (14) |
| Sp1413R | CATGCCTTCCAACAGCCG | (14) |
| lgt-CmF | TTATAAAAGCCAGTCATTAG | (14) |
| lgt-CmR | TTTGATTTTTAATGGATAATG | (14) |
| *For cloning pEC210, pEC211.* | | |
| BamHI lgt comp F | GATAGGATCCACGAACGACTGACAAG | This work |
| XhoI lgt comp R | GATACTCGAGACATTTAGTTTTCCTCCTCTG | This work |
| *Used to generate transforming DNA for making ECSPN211.* | | |
| 00322 up F | GCTTCAATGTACCACGAAG | This work |
| kan-00322 up R | ACGAACTCCAATTCACTGTTCAGCTTTACCATAATAAGACCTC | This work |
| kan-00322 down F | TCTGAAGTACATCCGCAACTAGACTAAATGGTAGCTCTCTG | This work |
| 00322 down R | CATCATTCATAAAATGGTCGTC | This work |
| pABG kanF | GAACAGTGAATTGGAGTTCG | This work |
| pABG kanR | AGTTGCGGATGTACTTCAG | This work |
| *For cloning pEC213.* | | |
| BamHI 00322 comp F2 | AGGTAGGATCCGTGGAAATGGTAATCACACTG | This work |
| XhoI 0322 comp R | GGTTATCTCGAGGGCAGTAGTAGCTCTCTG | This work |
